# CC-Tempo: A cell-cell communication aware temporal model of cellular dynamics

**DOI:** 10.1101/2023.12.04.569835

**Authors:** Sheikh Saifur Rahman Jony, M. Sohel Rahman, Md. Abul Hassan Samee

## Abstract

Delineating the mechanisms underlying cell state changes is key to gaining insights into organismal development and disease prognosis. Various methods have been proposed to study cellular differentiation and cell fate specification. However, they either do not incorporate temporal information or do not consider the vital role of intercellular communication in cellular differentiation and cell fate determination. Furthermore, many of these methods lack interpretability, making it difficult to identify the critical genes and pathways that influence the differentiation process. Here we propose CC-Tempo, a cell-cell communication-aware model of cellular dynamics that leverages intercellular communication scores and can help identify important genes and pathways crucial for different stages of differentiation in various lineages. While previous studies have indicated that scRNA-seq data alone may not suffice for accurately predicting cell fates, CC-Tempo demonstrates that incorporating intercellular communication significantly enhances the performance of such models. CC-Tempo can predict the significance of genes and pathways at different stages of the differentiation process. By perturbing these genes in silico, CC-Tempo reveals their efficacy for manipulating cell fate, which can be crucial for defining efficient reprogramming factors.

## Introduction

In multicellular organisms, life begins with a single zygote cell that gives rise to a complex organism comprising specialized cell states. The process determining such specialization is known as cellular differentiation, a cornerstone topic in developmental biology^1^. Researchers actively study this field to answer key questions, such as tracing a cell’s differentiation history, understanding the transcriptional regulatory mechanisms governing differentiation, and predicting its transcriptional state (“cell fate”) at future time points. Traditional bulk data fall short in answering these questions as they cannot capture variations between individual cells or track them over time. Single-cell RNA sequencing (scRNA-seq) has the potential to address these limitations by providing transcriptomic data at the single-cell level^2^. However, scRNA-seq technologies necessitate cell death for data collection, making it impossible to track a cell over time. Recent computational models of temporally resolved scRNA-seq data have shown promising advances to mitigate this issue, although there remain critical gaps in building mechanistic models that elicit actionable hypotheses. In this manuscript, we propose CC-Tempo (A cell-cell communication aware temporal model of cellular dynamics) to address this gap.

A large number of methods have applied “pseudo-time ordering” to study cell state transitions in scRNA-seq data. These methods order cells along pseudo-temporal axes. The inferred pseudo-temporal values have been employed to forecast cellular trajectories and, ultimately, cell fate^3,4^. However, for temporally-resolved scRNA-seq data, these pseudo-time ordering methods are unable to take advantage of the information on distinct time points^5,6^. In order to model cellular differentiation and cell state changes using temporally-resolved data, several new approaches have emerged^7–10^. Among them, some attempted to model differentiation as a process but faced limitations in their solving or modeling capabilities^9^. There exist other methods as well, mostly centered around the concept of optimal transport (OT)^11^. These methods essentially construct maps between the scRNA-seq data of consecutive time points to form a continuum of cell states akin to Waddington’s metaphorical epigenetic landscape^12,13^. Schiebinger et al.^8^ proposed a framework aimed at learning this metaphorical epigenetic landscape from scRNA-seq data. Their approach is grounded in the concept that cells at specific time points are drawn from a distribution within the gene expression space, encompassing both ancestral and descendant cellular distributions. This underlying distribution was estimated using an unbalanced OT formulation. Overall, these methods have set a strong rationale that computational modeling can elicit a deeper understanding of cellular differentiation by leveraging temporally-resolved scRNA-seq data. Further opportunities for rigorously building these models are offered by the cutting-edge LT-scSeq technology (lineage tracing with single-cell RNA sequencing)^14,15^. In LT-scSeq, cells are tagged during initial time points to subsequently track their descendant cells at later stages. LT-scSeq data combines lineage tracing information with scRNA-seq measurements and are considered gold standards for benchmarking the aforementioned computational models^16^.

Weinreb et al.^17^ featured an LT-scSeq dataset of mouse hematopoiesis. They evaluated various existing methods for predicting cell fate based on single-cell RNA sequencing (scRNA-seq) data, including Population Balance Analysis (PBA)^9^, WaddingtonOT^8^, and FateID^7^. Importantly, they concluded that single-cell gene expressions alone are insufficient for accurately predicting cell fate outcomes. Following up, Yeo et al.^18^ proposed a model named PRESCIENT that takes cell proliferation into account and outperformed the above methods on Weinreb et al.’s dataset. Introducing the notion of “potentials” within the space of transcriptional cell states, PRESCIENT viewed cellular differentiation as a change of potentials over time, which can be described by a diffusion equation. In contrast to the other methods, PRESCIENT is a generative model, and Yeo et al. showed interesting applications of PRESCIENT on unobserved data points and to predict cell fates under transcription factor perturbations.

Notably, all existing methods model cell state transitions by considering each cell as an independent, isolated entity. Yet, cells do not exist in isolation within an organism; they coexist within tissues and engage in cell-cell interaction. Indeed, Schiebinger et al.^8^ discussed that the assumption of a cell’s autonomous trajectory is likely inaccurate. It is well understood that cells attain distinct functional states not only through their individual gene expressions but also through local cell-cell communication^19^. This intercellular crosstalk, often facilitated by ligand-receptor pairs, is crucial for steering diverse cellular decisions, spanning from cell cycle regulation, cell death, and migration to lineage-specific differentiation ^20–22^. Thus, we propose CC-Tempo, a cell-cell communication aware temporal model of cellular dynamics. Even though single-cell RNA sequencing (scRNA-seq) data doesn’t explicitly offer insights into cell-to-cell communication, it inherently carries details about ligand-receptor interactions, which can be harnessed to deduce intercellular communication potentials ^23,24^. CC-Tempo demonstrates that incorporating these intercellular interactions makes it possible to more accurately capture the underlying dynamics of cell trajectories compared to existing methods. A second notable feature of CC-Tempo is its easy and immediate interpretability.

## Results

### CC-Tempo: A cell-cell communication aware temporal model of cellular dynamics

CC-Tempo defines the temporal changes in cells as a combination of two interrelated diffusion processes (See Sec. 4 (Methods)). The first one is a diffusion process over the single-cell RNA expression space, which is modeled by a linear potential function (Eq. 1) following previous studies^8,18^. This linear potential function is learned from the temporal snapshots of single-cell RNA expression data using a feed-forward neural network. However, incorporating intercellular communication into the model is quite tricky as it involves the intricate communication of multiple cells. Moreover, existing methods^19,25–27^ can only infer intercellular communication for different cell types present in the dataset rather than at single-cell resolution, which further complicates the already intricate issue.

In tissues, intercellular communications are carried out via physical interactions, exchange metabolites, and ligand-receptor signaling^20^. Ligand-receptor-mediated signaling has been widely studied in the literature, and comprehensive databases of ligand-receptors exist that can characterize most of the intercellular communication^28,29^. This ligand-receptor-mediated communication plays a critical role from early embryonic development to various tissue development to dysregulation like cancer, autoimmune and metabolic diseases.

Interestingly, the expression level of these ligand-receptors can be directly extracted from scRNA-seq data. Moreover, these pairs can be organized into groups called signaling pathways, offering a concise overview of intercellular communication patterns^30^. While existing methods can infer such pairs and enriched signaling pathways across various cell types, they lack the granularity to do so at the single-cell level. To overcome this issue, we develop a Bayesian approach that enables the calculation of intercellular communication at the single-cell resolution using the output of these methods at the cell type level.

We model the intercellular communication of a single-cell as all the enriched incoming and outgoing communications via various signaling pathways from that cell to all other cells. Each of these signaling pathways will comprise each dimension in our intercellular communication space. To score the incoming (outgoing) communications of a single-cell, we calculate the aggregated expression of ligands (receptors) for each signaling pathway. As we already have information on the cell type and cell type-specific intercellular communications derived from existing methods, which is our Bayesian prior, we use this prior to update the communication score for each single cell in a Bayesian fashion (Fig 1A). We concatenate the incoming and outgoing scores into a single vector, which will be the input to our intercellular communication component—a distinct diffusion process illustrated in Fig 1A (See Eq. 2).

**Figure 1:**
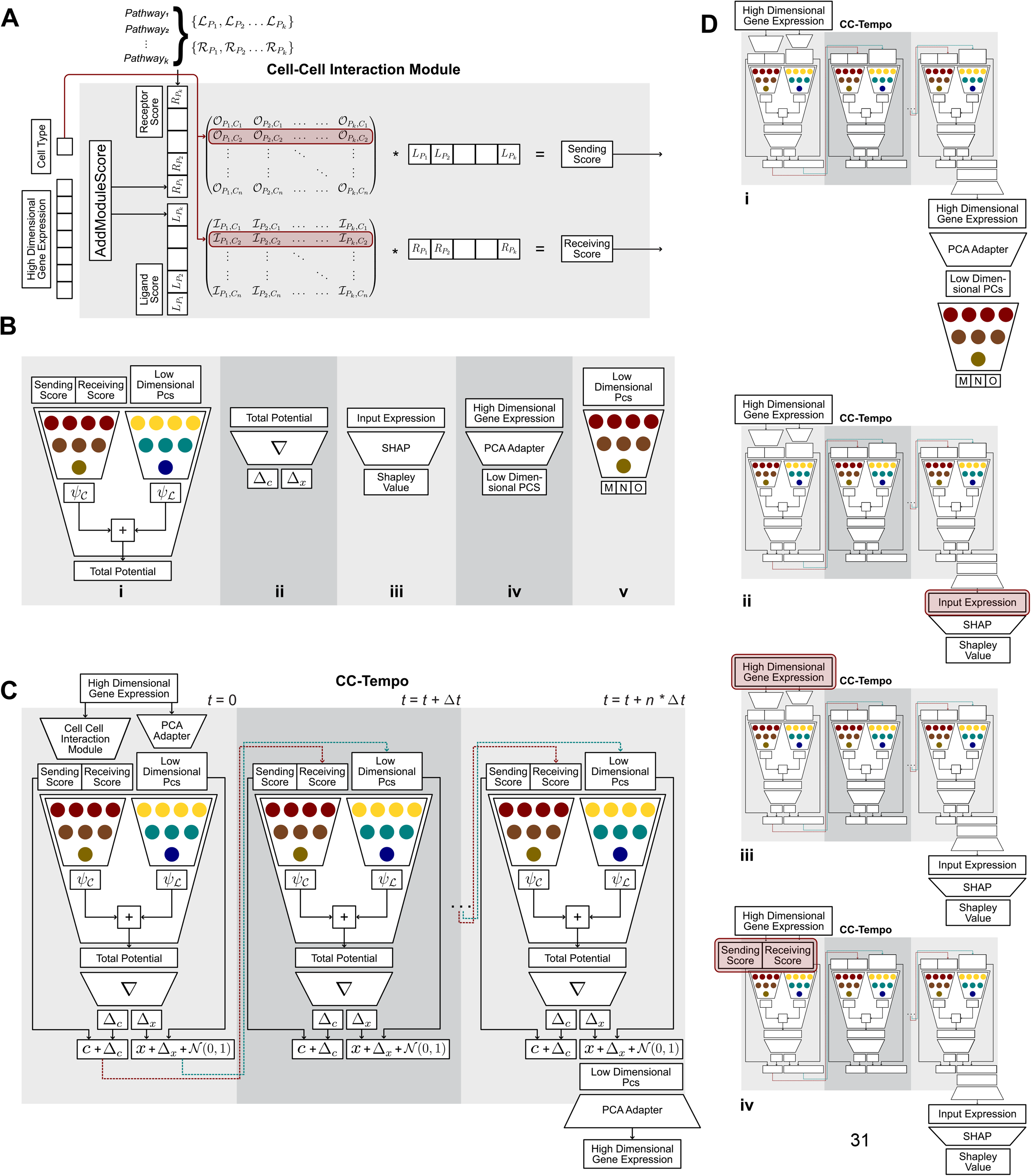
**A)** The cell-cell interaction module calculates the intercellular communication scores given scRNAseq gene expression data and cell type. It calculates the score using a set of signaling pathways that are enriched in the dataset and their incoming and outgoing score matrix. It calculates the Hadamard product of the corresponding row determined by the cell type with the corresponding ligand and receptor score of that cell to produce the sending and receiving score. B) An overview of all the components of CC-Tempo. **i)** The potential learning network and its components. It consists of two separate neural networks, of which the first takes intercellular communication score as input and outputs cell-cell interaction potential as output. In contrast, the second neural network takes the PCs of scRNA-seq gene expression data and outputs linear potential. Finally, the network sums both of the aforementioned potentials and outputs the total potential as its final output. **ii)** The gradient calculator - this component takes the total potential as input. It calculates the gradient of this potential with respect to both of the inputs, i.e., intercellular communication score and the PCs of scRNA-seq expression data. **iii)** SHAP is used to find out the importance of different input features for a given output/outputs and ranks the inputs based on shap score accordingly. **iv)** The PCA Adapter works in both directions, i.e., it can project the high dimensional input to low dimensional input or vice versa. It consists of a layer of neural networks. **v)** The cell type classifier takes the PCs of the scRNA-seq gene expression data and classifies that input into either monocytes, neutrophils, or other cell types. **C)** The full CC-Tempo model: It takes high dimensional gene expression of cells as an earlier time point, simulates the cells over the specified time points, and outputs the final day high dimensional gene expression of those cells. **D)** Different usage of CC-Tempo **i)** CC-Tempo can be used to simulate cells to later time points and classify them into different cell types. **ii)** CC-Tempo can be used to evaluate the gene expression of the final day and determine genes that are significant for determining final fates. **iii)** CC-Tempo can be used to simulate any intermediate time points and the genes that are important on those time points to determine the final day cell fates. **iv)** CC-Tempo can be used to evaluate the importance of signaling pathways in a similar fashion as well.

This second difussion process is also learned by a feed-forward neural network and the output of these two aforementioned networks are combined to derive next state of a cell (Fig 1B). Thus, to CC-Tempo, a cell state is a tuple comprising single-cell RNA expression data and intercellular communication score. Unlike existing methods ^8,9,18^, CC-Tempo learns this potential directly from the single-cell RNA expression data without reducing the data into a low dimensional space (Fig 1C, See Sec. 4 (Methods)). Utilizing the full gene expression data as CC-Tempo’s input allows it to be interpretable in various ways (Fig 1Di-iii).

Both components of the state evolve over time by their corresponding drift velocity, which takes incremental movement towards the direction of lower potential in their corresponding potential space (Fig 1C). Particularly, the drift velocity is the negative gradient of their corresponding potential value (Fig 1Bii). The linear potential drives the cells into lower potential space as the cells continue shifting to their final cell fate over time. The cell-cell interaction potential maps how cell communication changes with each other as they proceed to their final fate state. These two stochastic processes are simulated by their first-order time discretized equations to obtain the cell’s movement at each time step. These equations can be used iteratively to obtain the cell state at the final time point, which is then fitted with the observed cell state tuple using the objective function (Fig 1C). The gene expression loss is calculated by minimizing the Wassenstein loss between the scRNA-seq data and model-predicted gene expressions, while the intercellular communication score loss is a Wasserstein loss between intercellular communication scores estimated from the data and those predicted by the model. The final loss is the summation of these two losses and a regularization term. Wasserstein loss has been widely used in learning such a diffusion process^31,32^.

### CC-Tempo outperforms existing methods when applied to mouse hematopoiesis data

We apply CC-Tempo to Weinreb et al.’s mouse hematopoiesis LT-scSeq dataset^17^ comprising 130,887 cells of 11 cell types and collected at three time points: Day 2, Day 4, and Day 6. Furthermore, LT-scSeq provided clonal data for 49,302 out of the total cells. A “clone” refers to a group of cells originating from the same progenitors in previous time points. In total, there are 5,864 unique clones in this dataset (Fig. 2A, B). The lineage tracing data allows us to validate CC-Tempo’s performance on developmental cell differentiation and benchmark CC-Tempo with others. Specifically, following the dataset^17^ and subsequent methods^8,18^, we calculate the metric clonal fate bias to evaluate CC-Tempo’s predictions on cell fate specification. In the original dataset, clonal fate bias is defined as the total number of neutrophils divided by the total number of neutrophils and monocytes for that specific clone. So, a clonal fate bias closer to 0 denotes the fate of a progenitor cell to be a monocyte, while a clonal fate bias closer to 1 denotes the fate of a neutrophil (Fig. 2A, C). Following PRESCIENT^18^, we derive clonal fate bias from CC-Tempo by making 100 copies of progenitor cells and then simulating them till Day 6 via CC-Tempo to get the final state. Each of these simulated Day 6 cells will possess unique states as a result of the model’s random noise component. We classify the final state of each copy as neutrophil, monocyte, or other types via a simple Logistic Regression classifier to emphasize on the learning capability of CC-Tempo, rather than using sophisticated classfier to obtain better results.. Finally, following the original study^17^, we assess the performance of CC-Tempo by measuring the Pearson correlation coefficient (PCC) of the model predicted fate bias with respect to the actual clonal fate bias provided by the lineage tracing data. We benchmarked CC-Tempo on the aforementioned metrics against PRESCIENT^18^ since it was shown to outperform the other existing methods on Weinreb et al.’s dataset. These other methods are Population Balance Analysis (PBA), WaddingtonOT, and FateID. When our manuscript was in the editing stage, another relevant tool named TIGON was published by Sha et al.^33^. It is a theoretically appealing model that simultaneously infers cell velocity, growth, and dynamics. Similar to our CC-Tempo and Yeo et al.’s PRESCIENT, Sha et al. evaluated TIGON on Weinreb et al.’s lineage tracing data. Both of the models had competitive performance, but PRESCIENT had a simpler network. Hence, we were attracted to it. CC-Tempo outperformed PRESCIENT in both Pearson correlation coefficients (PCC). Compared to PRESCIENT, CC-Tempo achieved a 4% improvement in PCC (improved to 0.52 from 0.50) (Fig. 2D). To confirm that the intercellular communication actually playing some role in model’s output, we dropped top 5 and then top 10 signaling pathways ranked by CC-Tempo (See Methods). Dropping top 5 signaling pathways decreased the PCC to 0.512 and dropping the top 10 signaling pathways declined the PCC to 0.50, signifying that intercellular communication has helped CC-Tempo capture cell fate more accurately. We trained all these above settings 100 times and evaluated the above score to test the statistical significance of the improvement (Supplementary Table 1). CC-Tempo clearly outperformed PRESCIENT (p-value 7e-19) and restricted version of CC-Tempo when top 10 signaling pathways were dropped (p-value 7e-5). However, CC-Tempo and CC-Tempo when top 5 signalling pathways were dropped do not have any significant performance difference (p-value 0.09). Additionally, we calculate clonal fate bias deviation (See Sec. 4 (Methods)) and draw the distribution plot of this deviation (Supplementary Figure 1A). As is evident from the figure, CC-Tempo has a much shorter tail than PRESCIENT, signifying that CC-Tempo tends to stick closer to actual clonal fate bias than PRESCIENT.

**Figure 2:**
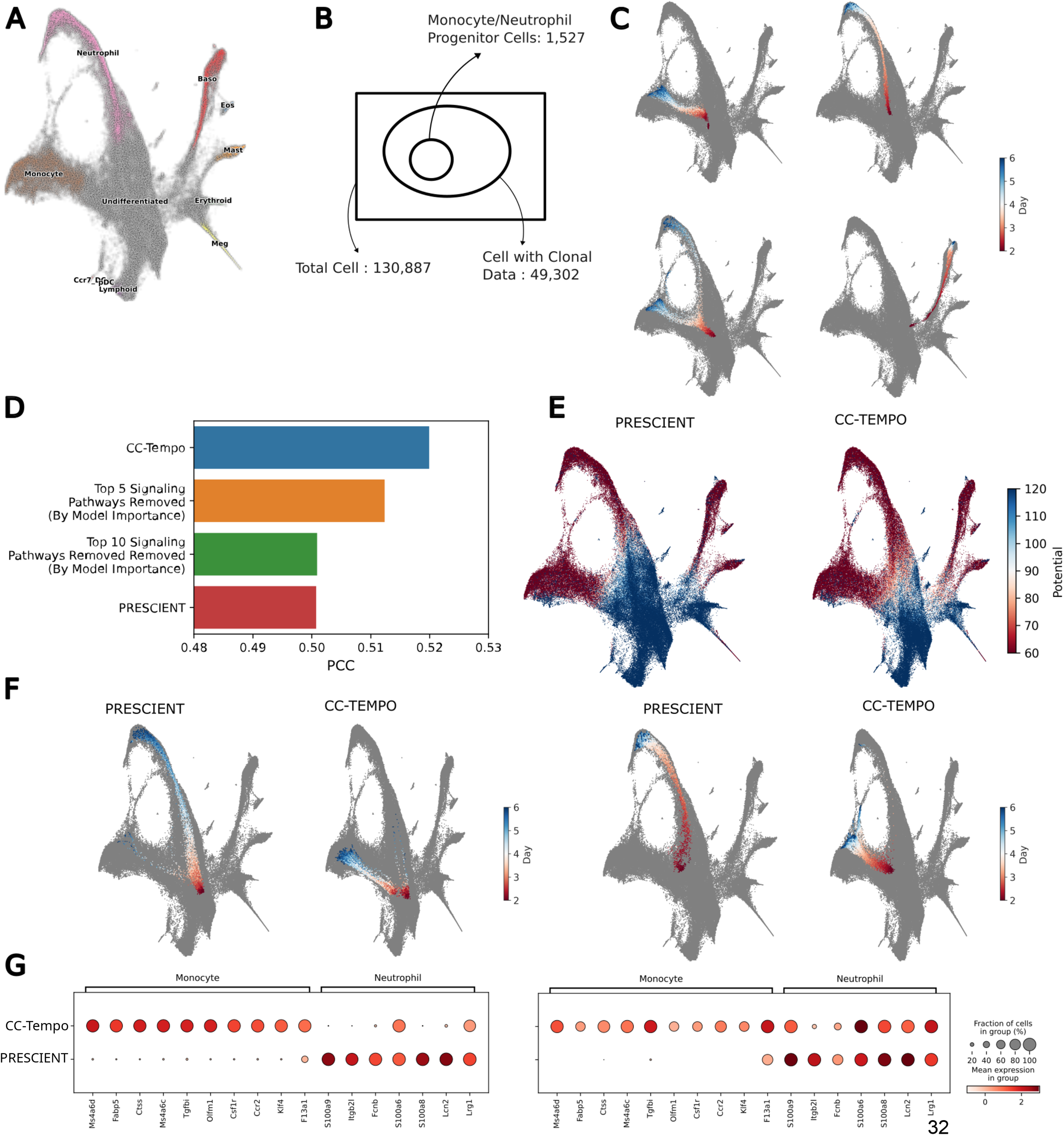
**A)** UMAP of Weinreb et al.’s mouse hematopoiesis LT-scSeq dataset. The dataset contains 11 different cell types. **B)** A quick overview of the number of cells in total in the dataset, the number of cells having clonal information (from Lineage Tracing), and the number of cells that are progenitors on Day 2 for monocytes or neutrophils. **C)** Examples of different clonal fate biases. (Top-Left) A group of trajectories having very low (∼0) clonal fate bias. This fate corresponds to monocytes and went into the monocyte branch. (Top-Right) A group of trajectories having very high (∼1) clonal fate bias. This fate corresponds to neutrophil went into the neutrophil branch. (Bottom-Left) A group of trajectories entering both of the possible paths and having a possible 0.5 clonal fate bias. (Bottom-Right) A group of trajectories entering none of the monocyte or neutrophil paths and hence resulting in a 0.5 clonal fate bias solely due to pseudo-count. **D)** Plots showing the comparison of PCC scores of CC-Tempo trained on different sets of intercellular communication pathways and a state-of-the-art model, PRESCIENT. **E)** The total potential of different cells in the UMAP predicted by PRESCIENT (Left) and CC-Tempo (Right). **F)** Two random examples of trajectories where CC-Tempo predicts the correct trajectories, hence correct fate bias, while PRESCIENT fails to do so. **G)** The expression of marker genes of monocytes or neutrophils in the final day cells by CC-Tempo and PRESCIENT for the previous two examples.

Furthermore, while comparing the potential value of CC-Tempo and PRESCIENT, we find that cells in PRESCIENT tend to stay in high potential even after the bifurcation of the monocyte and neutrophil in UMAP, but cells in CC-Tempo tend to go into lower potential at the bifurcation points which suggests that CC-Tempo tends to capture cell fate information much earlier in time than PRESCIENT (Fig. 2E). We have plotted some examples of the cell trajectory where PRESCIENT fails to capture the clonal fate bias while CC-Tempo accurately predicts the score (Fig. 2F). Overall, this benchmarking set a strong premise for applying CC-Tempo to define the genes and intercellular communications underlying cell state specification during hematopoiesis.

### CC-Tempo accurately predicts marker gene expression in Day 6 cells and identifies the cell-fate-defining genes

We next investigated why CC-Tempo outperforms PRESCIENT. We posited that a method’s performance in predicting cell fate bias depends on its accuracy in predicting marker gene expression in Day 6 cells. Thus, we used both methods to predict marker gene expressions in Day 6 cells. Checking these details is straightforward for CC-Tempo since it directly works with high-dimensional gene expression data and outputs the final cell’s gene expression as well as intercellular communication score in the same high-dimensional gene space. However, many models, including PRESCIENT, use low dimensional representation to train their model, hence making it complicated to interpret the model. The advantage of CC-Tempo is that, it incorporates the dimensionality reduction (expansion) as a part of the Neural Network (See Methods), which allows us to individual genes as inputs to CC-Tempo and leverage different model interpretability techniques to assign predictive importance to each gene. The previous methods use reduced dimensional representation of gene expression data as inputs, which limits model interpretation techniques to prioritize only the important dimensions, but not individual genes..

Following the dataset^17^ and subsequent methods^8,18^, we evaluated CC-Tempo on Neutrophil vs Monocyte cell fate specifications. We obtained the monocyte and neutrophil marker gene annotations from Weinreb et al.’s original study (Supplementary Table 2). We visualized their expression as predicted by CC-Tempo and PRESCIENT in Day 6 cells. We focused on the cells where CC-Tempo’s predictions are more accurate than PRESCIENT’s in cell fate specification(Fig. 2G). This comparison indicates that a model’s accuracy is highly dependent on its ability to predict marker gene expression. As an orthogonal approach to defining the genes driving CC-Tempo’s accurate predictions, we applied SHAP (SHapley Additive exPlanations) on the fates predicted by CC-Tempo for Day 6 cell. SHAP (SHapley Additive exPlanations) is a widely used local model explanation tool for neural networks^34–36^. For a given input, it identifies the most influential variables in determining the output and ranks them according to their importance. Here, we used SHAP to identify and rank the genes driving CC-Tempo’s classification accuracy for Day 6 cells. Briefly, we take all the progenitor cells (Day 2), simulate them via CC-Tempo, and derive their Day 6 (final day) cell state. We then use our trained logistic regression classifier to classify these predicted cells into neutrophils, monocytes, or other cells. We pass the predicted monocyte cells’ gene expression to obtain the linear potential and explain this with SHAP to determine which genes were highly important for that potential. Interestingly, the top 15 most important genes for the monocyte lineage showed one out of ten of the previous monocyte markers reported by Weinreb et al. (*Fabp5*; Fig. 3A). However, 14 of these 15 genes are indeed significantly differentially expressed between monocytes and all other cell types, as we confirmed using the Wilcoxon’s rank-sum test (Fig. 3C, Supplementary Table 3).

**Figure 3:**
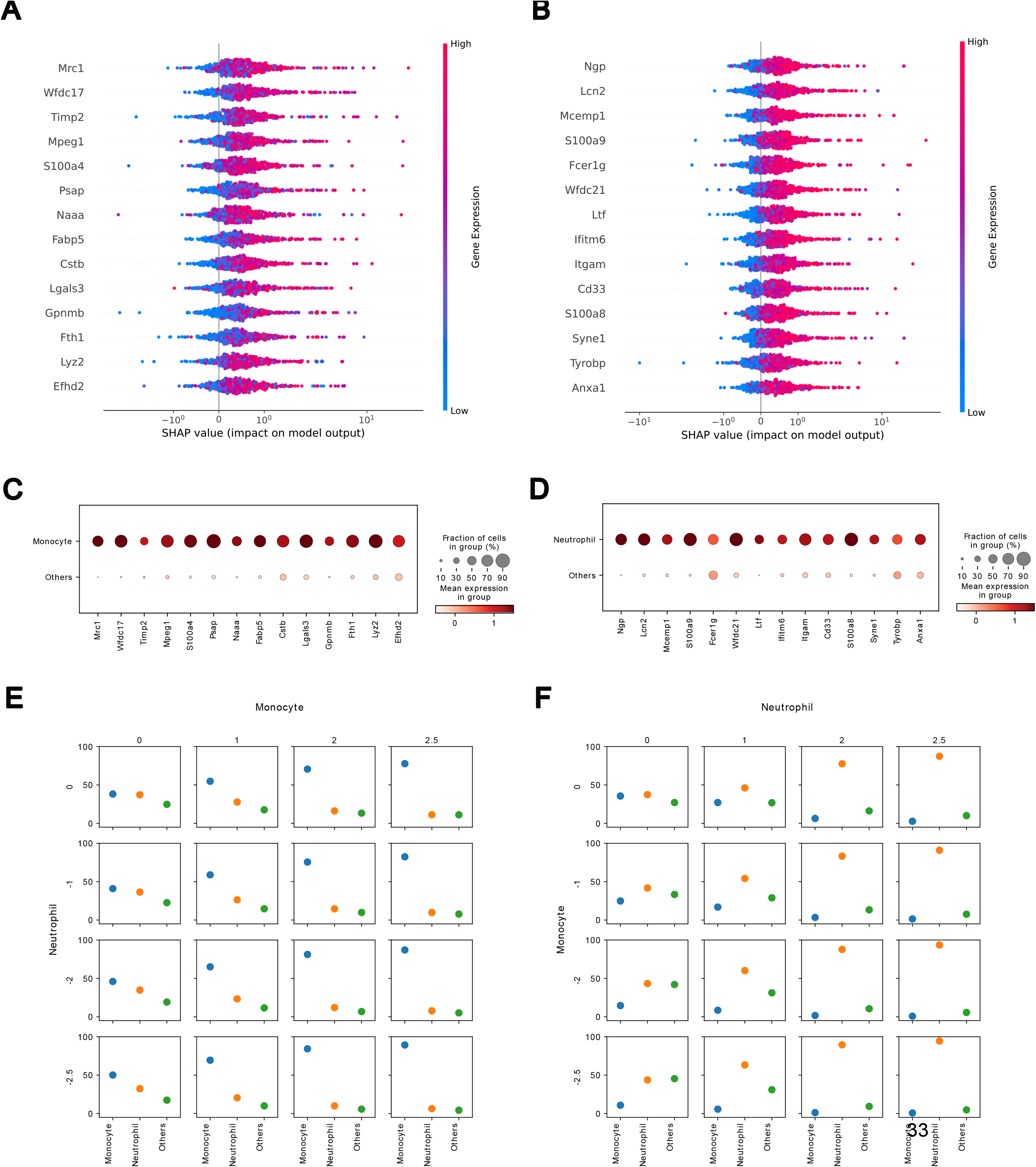
Beeswarm plot showing the most important genes for determining final day cell fate for **A)** monocyte lineages and **B)** neutrophil lineages along with gene expression. **C)** The dot plot of gene expression of these important genes for monocyte cells and all other cells from the dataset on the final day (Day 6). **D)** The dot plot of gene expression of these important genes for neutrophil cells and all other cells from the dataset on the final day (Day 6). **E)** A plot of the percentage of cells in different lineages (monocytes, neutrophils, and others) when different transcription factors of monocytes are gradually upregulated while the transcription factors of neutrophils are downregulated. **F)** A plot of the percentage of cells in different lineages (monocytes, neutrophils, and others) when different transcription factors of monocytes are gradually downregulated while the transcription factors of neutrophils are upregulated.

Additionally, we find a new gene *1110002J07Rik*, suggested by SHAP. However, Wilcoxon’s rank-sum test shows that it is not significantly differentially expressed between monocytes and all other cell types. (Supplementary Figure 1B).

Following the similar method above, we derive the most important genes for neutrophil lineage. In a similar fashion, the top 15 most important genes for the neutrophil lineage showed three of the previous neutrophil markers reported by Weinreb et al. (*Lcn2, S100a9, S100a8*; Fig. 3B). Moreover, 14 of these 15 genes are indeed significantly differentially expressed between neutrophils and all other cell types, as we confirmed using the Wilcoxon’s rank-sum test (Fig. 3D, Supplementary Table 4). Additionally, we find that *1110002J07Rik* plays an important role in neutrophils’ fate determination as well, according to SHAP. Again, it is not significantly differentially expressed between neutrophils and all other cell types.

### CC-Tempo predicts expected fate changes in cells with progenitor transcription factors’ perturbation

By design, CC-Tempo can predict the differentiation trajectory starting from any cell state. Thus, as long as it captures the mechanisms underlying cellular differentiation dynamics from the observed data, CC-Tempo can reliably predict the outcome of genetic perturbations (See Sec. 4 (Methods)). We next utilized CC-Tempo to perturb transcription factor (TF) expression in early progenitor cells and investigate how that affects the final day cell fate.

In particular, as above, we choose the Day 2 progenitor cells with at least one monocyte or neutrophil in their clone on the final day and perturb the corresponding TFs for neutrophils or monocytes to check if CC-Tempo predicts a fate bias change toward the correct fate. A comprehensive list of TFs expressed in monocytes and neutrophil progenitor cells is available in the literature^37–41^. The following transcription factors are available in Weinreb et al. dataset for monocytes *F13a1, Ms4a6c, Ly6c2, S100a4, Rassf4, Csf1r, Hpse, Ly86, Emb, Papss2, Ctss, Slpi, Irf8, Nr4a1*, and *Klf4*. The transcription factors for neutrophils that are available are *Ltf, Ngp, Lcn2, Cd177, Camp, S100a9, Ifitm6, Itgb2l, Pglyrp1, S100a8, Lrg1, Fcnb, Gp1bb, Lyz2*, and *Syne1*.

First, we derive the percentage of different types of cells in final fate when none of the transcription factors are perturbed. Then we upregulate or knock down different transcription factors following the methods proposed by Yeo et al. They set the scaled normalized z-score expression value of target genes to less than 0 for knockdowns and greater than 0 for upregulation. For our experiments, holding the neutrophil transcription factors the same, we upregulate monocytes’ transcription factors gradually from unperturbed to 2. 5. This results in the increase of monocytes’ percentage in the final cell states from 35. 59 ± 0. 11 while unperturbed to 77. 72 ± 0. 09 while monocyte transcription factors were set 2. 5 (Fig. 3E). Similarly, we upregulate neutrophils transcription factors gradually from unperturbed to 2. 5 while holding the monocyte transcription factors the same. This increases neutrophil percentage in the final cell state from 37. 4 ± 0. 12 the unperturbed setting to 87. 30 ± 0. 06 (Fig. 3F).

Secondly, while holding the monocyte transcription factors constant, we gradually knock down the neutrophil transcription factors from an unperturbed state to −2.5. We see that the percentage of neutrophils decreases from 37. 4 ± 0. 12 to 32. 45 ± 0. 09 (Fig. 3E) while the percentage of monocytes increases from 35. 59 ± 0. 11 to 50. 15 ± 0. 11. In a similar manner, we hold the neutrophil transcription factors constant while we knock down the monocyte transcription factors from unperturbed to −2.5. This decreases monocyte from 35. 59 ± 0. 11 to 10. 80 ± 0. 08. On the other hand, the percentage of neutrophils increases from 37. 4 ± 0. 12 to 43. 77 ± 0. 11.

Finally, totally knocking down the neutrophils’ transcription factors (−2.5) and totally upregulating the monocyte transcription factors (2.5) results in the percentage of monocyte of 89. 31 ± 0. 07 and the percentage of neutrophils to be 6. 39 ± 0. 05. While totally knocking down the monocyte transcription factors and totally upregulating the neutrophil transcription factors results in the percentage of neutrophil being 94. 49 ± 0. 05 and the percentage of monocyte being 0. 71 ± 0. 01.

### CC-Tempo identifies genes whose activation at intermediate time points determines the final cell fate

We next applied CC-Tempo to investigate cellular differentiation mechanisms at intermediate time points. Such analyses can detect a “window of opportunity” and the most promising target genes (signaling pathways) for effective perturbation. We again used SHAP to find the initial day progenitor cells’ gene expressions that are imperative to determining final cell fate. CC-Tempo finds us that the progenitor cells’ top 10 genes that determine the monocyte fate are *Igfbp4, Casp6, Samhd1, Ifi203, Prdx1, Muc13, Rbms1, Spint2, Srgn, Tpd52* (Fig. 4A). On the other hand, the top 10 genes for neutrophil progenitor cells are *Srgn, Tmed3, Calr, Gstm1, Igfbp4, Dstn, Muc13, Spint2, Prdx1, Ifi203* (Fig. 4B).

**Figure 4:**
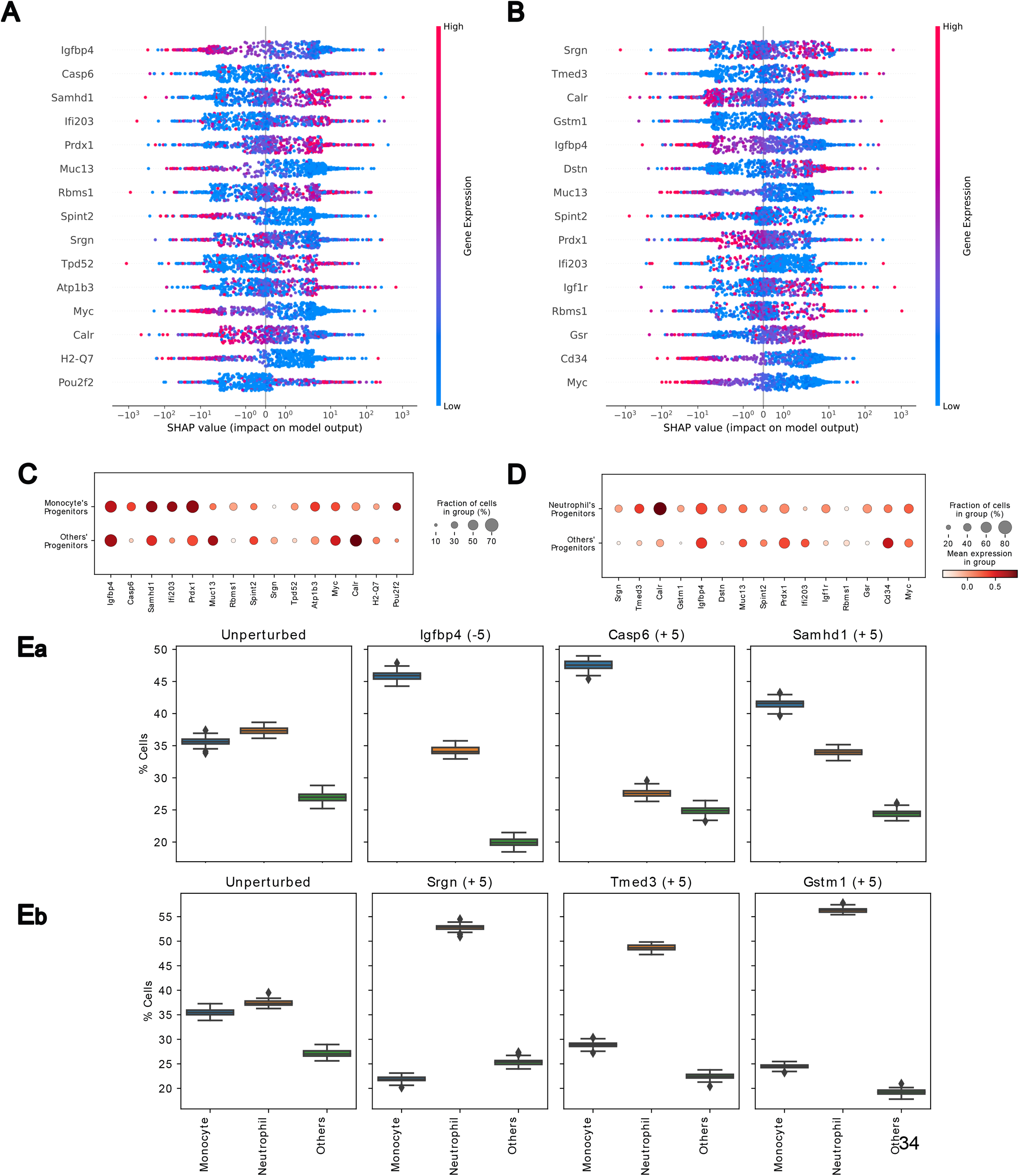
Beeswarm plot showing the most important genes in progenitor cells on Day 2 for determining final day cell fate for **A)** monocyte lineages and **B)** neutrophil lineages along with gene expression. **C)** The dot plot of gene expression of these important genes of progenitor cells for monocyte cells and all other cells. **D)** The dot plot of gene expression of these important genes of progenitor cells for neutrophil cells and all other cells. **Ea)** A plot of the percentage of cells in different lineages (monocytes, neutrophils, and others) when some of these important genes for monocyte lineage are perturbed in silico. **Eb)** A plot of the percentage of cells in different lineages (monocytes, neutrophils, and others) when some of these important genes for neutrophil lineage are perturbed in silico.

We further divided the progenitor cells into monocyte progenitor cells and other progenitor cells and plotted the gene dot plot for these important genes to check if they are actually differentially expressed between the aforementioned two types of progenitor cells (Fig 4C). The dot plot shows the significant distinction in many genes between the two types of progenitor cells. We further validated if these genes are indeed significantly differentially expressed between monocytes’ progenitors and all other cell types progenitors using Wilcoxon’s rank-sum test (Supplementary Table 5). The test yields that, indeed, all of the 15 genes are significantly differentially expressed between the two progenitor types. In a similar fashion, we did the same for neutrophils’ progenitor cells vs. all other types of cells and plotted their gene dot plot (Fig. 4D). We further calculated Wilcoxon’s rank-sum test to check if these genes are actually significantly differentially expressed between the two progenitor types. The test yielded 13 out of the 15 genes are significantly expressed between these two types of progenitors. Only *Igfbp4* and *Rbms1* were not significantly expressed (Supplementary Table 6).

Interestingly, not the high expression of all of these genes is helpful for a particular lineage; rather, some genes’s low expression can be important for final fate determination. For example, we can see that *Igfbp4* is an important gene for monocyte lineage, but the low expression of the gene is required to play a role in monocyte production. The same can be seen for *Muc13, Spint2*, and some other cells. The gene dot plot also shows a similar result. This phenomenon is also present in neutrophil progenitor cells where the low expression for *Muc13, Ifi203, Cd34*, and some other genes are important in neutrophil production. Moreover, the high expression of *Calr* is important for neutrophil lineage, while the low expression is important for monocytes. The same can be said for *Ifi203* and *Prdx1*

Next, we perturb some of these important genes in the progenitor cells and check if they can actually change the fate of the final day cells. We posited that if the expression of these genes is important for the final day’s fate, then their expression at the initial time point will have a significant effect on the fate of the final day cells, and changing their expression will change the cells’ fate. First, we calculated the percentage of different cell types at an unperturbed setting and found out that the percentage of monocytes is 35. 62 ± 0. 12. If we set the expression of *Igfbp4* to −5 and keep the expression of all other genes the same, we see that the percentage of monocytes increases to 45. 85 ± 0. 12. Similarly, setting the expression of *Casp6* and *Samhd1* to +5 increases the percentage of monocytes to 47. 58 ± 0. 13 and 41. 49 ± 0. 12 respectively (Fig 4Ea). We perturbed some of the important genes for neutrophils in a similar fashion as well. The percentage of neutrophils in an unperturbed setting was 37. 45 ± 0. 11. We set the expression of *Srgn, Tmed3*, and *Gstm1* to +5 separately, and this increased the percentage of neutrophils to 52. 71 ± 0. 08, 48. 76 ± 0. 11 and 56. 45 ± 0. 11 respectively (Fig 4Eb).

CC-Tempo can be used to determine important genes for any lineage at any intermediate time points rather than just the initial and final time points. Weinreb et al. dataset contain three time points: Day 2, Day 4, and Day 6. CC-Tempo can determine genes that are important at the intermediate time point, Day 4, for determining the final day fate of a cell. We first take all the monocyte or neutrophil progenitor cells as previously determined. We then simulate these cells to Day 4 via CC-Tempo. Then, again, we use SHAP to interpret the model. This time, we pass the Day 4 cells’ gene expression as input to CC-Tempo, simulate it till the final day, and determine the potential. Then, we use SHAP to explain which genes of Day 4 are highly influential in determining the final day’s fate.

We carry a similar analysis as we did for Day 2 and find out the genes that are important on Day 4 for the final day (Day 6) fate determination. The important genes at Day 4 for monocytes and neutrophils are determined using SHAP in a similar fashion (Fig. 5A, B). The corresponding gene dot plots are also shown, and they again show significantly differential expression for most of the genes (Fig. 5C, D). We further carried out Wilcoxon’s rank-sum test to check if these genes are actually significantly differentially expressed between the two cell types. The test yielded all of the 20 genes are significantly expressed between monocyte and other cell types cell while 17 out of 20 genes are significantly expressed between neutrophil and other cell types (Supplementary Table 7,8). It also shows that some of the marker genes of monocytes and neutrophils start to differentially express and also play a significant role as early as Day 4. For example, the monocyte marker *Ctss* is an important gene for monocyte lineage on Day 4, while the neutrophil markers *S100a9, S100a8*, and *Lcn2* are important genes for neutrophil lineage on the same day.

**Figure 5:**
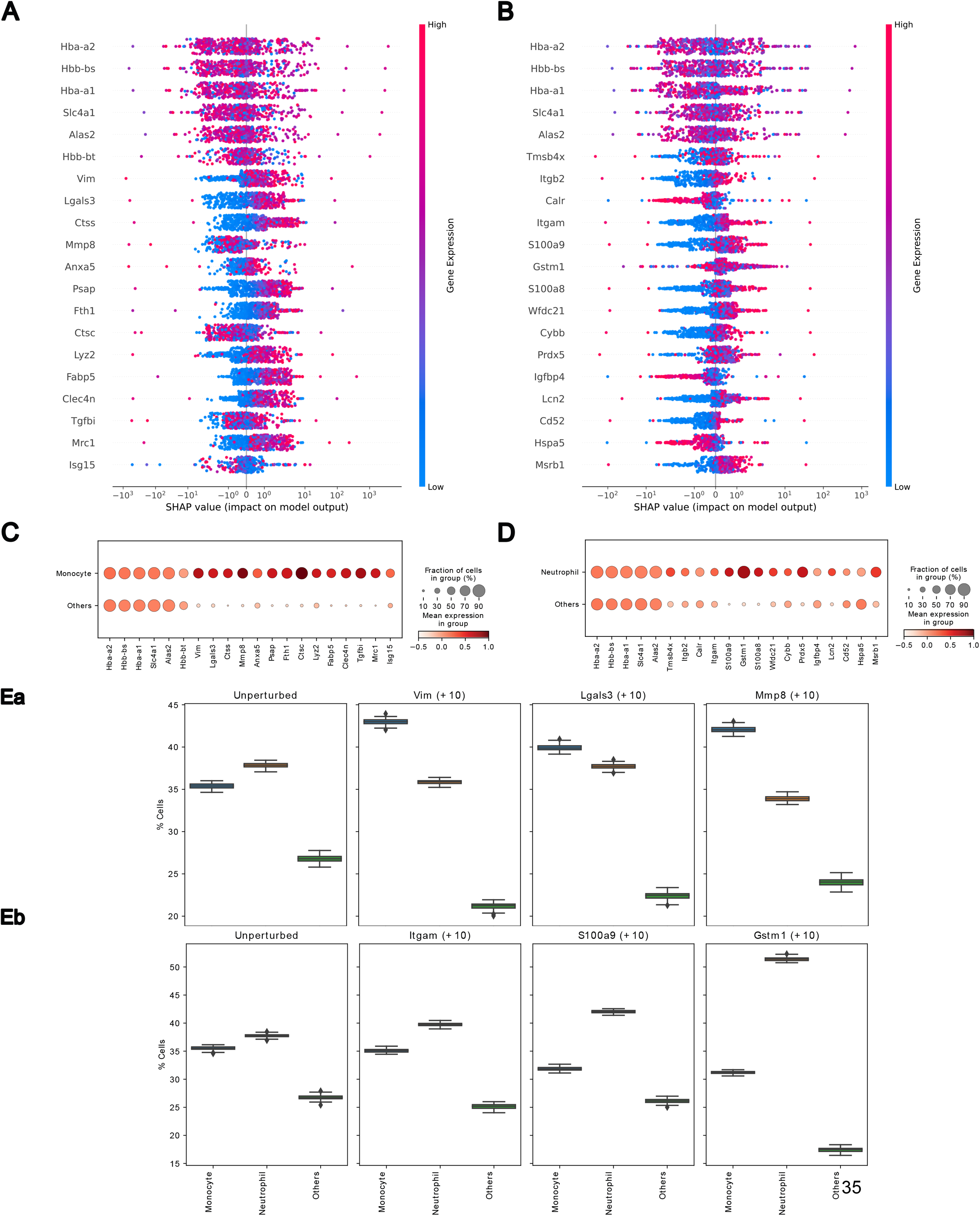
Beeswarm plot showing the most important genes on intermediate time point (Day 4) for determining final day cell fate for **A)** monocyte lineages and **B)** neutrophil lineages along with gene expression. **C)** The dot plot of gene expression of these important genes on intermediate time point (Day 4) for monocyte cells and all other cells. **D)** The dot plot of gene expression of these important genes on the intermediate time point (Day 4) for neutrophil cells and all other cells. **Ea)** A plot of the percentage of cells in different lineages (monocytes, neutrophils, and others) when some of these important genes for monocyte lineage are perturbed in silico. **Eb)** A plot of the percentage of cells in different lineages (monocytes, neutrophils, and others) when some of these important genes for neutrophil lineage are perturbed in silico.

In a similar fashion as above, we perturb some of these important genes in the Day 4 cells and check if they can actually change the fate of the final day cells. As we progress further in time, the perturbation of different genes tends to change the cell fate in a smaller amount. So, we set the value of the gene to +10/-10 to see if these important genes on Day 4 can actually change the final fate. First, we calculated the percentage of different cell types at an unperturbed setting and found out that the percentage of monocytes is 35. 39 ± 0. 06. If we set the expression of *Vim* to +10 and keep the expression of all other genes the same, we see that the percentage of monocytes increases to 42. 97 ± 0. 07.

Similarly, setting the expression of *Lgals3* and *Mmp8* to +10 increases the percentage of monocytes to 39. 89 ± 0. 07 and 42. 07 ± 0. 07 respectively (Fig 5Ea). We perturbed some of the important genes for neutrophils in a similar fashion as well. The percentage of neutrophils in an unperturbed setting was 37. 77 ± 0. 05. We set the expression of *Itgam, S100a9*, and *Gstm1* to +10 separately, and this increased the percentage of neutrophils to 39. 77 ± 0. 06, 41. 96 ± 0. 05 and 51. 33 ± 0. 05 respectively (Fig 5Eb).

We find that *Anxa2, Sirpa, S100a4, Igfbp4, Casp6*, and *Myc* are important genes throughout all time points in determining the fate of monocytes. Among them, except *Anxa2*, all other genes are differentially expressed among monocytes and all other cell types at each of the time points (Supplementary Table 9). We find out that the expression of *Anxa2, Sirpa, S100a4, and Casp6* keeps increasing for monocytes with time while the expression for other cell types remains almost the same or keeps decreasing. On the other hand, the expression of *Igfbp4* and *Myc* keeps decreasing over time for monocytes (Supplementary Figure 1C). Similarly, we find that the expression of *Msrb1* keeps increasing for neutrophils with time but not for other cell types, but the expression of *Eef1a1* keeps decreasing over time (Supplementary Figure 1D). All genes are differentially expressed among neutrophils and all other cell types at each of the time points (Supplementary Table 10).

Furthermore, CC-Tempo can be used to find important genes in unseen time points, for example, Day 3 or Day 5 as well in a similar fashion.

### CC-Tempo’s interpretation strongly agrees with experimental literature

CC-Tempo predicts that the expression of *Igfbp4, Casp6, Samhd1, Ifi203, Prdx1, Muc13, Rbms1, Spint2, Srgn, Tpd52, Atp1b3, Myc, Calr, H2-Q7* and *Pou2f2* are important in progenitors cell for monocyte differentiation while the expression of *Srgn, Tmed3, Calr, Gstm1, Igfbp4, Dstn, Muc13, Spint2, Prdx1, Ifi203, Igf1r, Rbms1, Gsr, Cd34* and *Myc* are important in progenitors for neutrophil differentiation (Fig. 4A-B). Next, we investigate the existing literature to inspect these genes contribution in monocyte vs neutrophil differentiation.

Weinreb et al. provide a list of genes whose expression is pivotal for determining cell fate in their dataset ^17^. According to CC-Tempo, *Igfbp4, Rbms1, Srgn, Myc, Muc13*, and *Calr* play important roles in both monocyte and neutrophil differentiation. Notably, Weinreb et al. identify *Igfbp4, Srgn, Myc, Muc13*, and *Calr* as crucial for neutrophil differentiation, while *Rbms1* is critical for monocyte differentiation. *Casp6, Samhd1*, and *Pou2f2* are essential only for monocyte differentiation, whereas *Gstm1, Igf1r*, and *Gsr* are crucial only for neutrophil differentiation. These findings are supported by the list provided by Weinreb et al. Additionally, the persistent expression of the interferon-stimulated gene *Ifi203* in monocyte progenitor cells has been reported ^42^. Literature also highlights that *Prdx1* and *Atp1b3* are important for monocyte/macrophage differentiation ^43,44^. *Spint2*, reported as essential for both monocyte and neutrophil differentiation, plays a key role in mediating macrophage recruitment and polarization^45^ while *Tmed* is important for neutrophil recruitment ^46^. Giladi et al. ^38^ also report the *Cd34* gene as significant for neutrophil differentiation. Lastly, *H2-Q7* is identified as crucial for basophil progenitors by Weinreb et al., whereas *Tpd52* and Dstn have no known role in monocyte or neutrophil differentiation yet.

CC-Tempo also provides a comprehensive list of genes that are crucial at intermediate time points for monocyte neutrophil differentiation (Fig. 5A-B). CC-Tempo reports that haemoglobin genes such as, *Hba-a2, Hbb-bs, Hba-a1,Hbb-bt* and *Slc4a1*, are important for both monocyte and neutrophil differentiation. It has been reported that the protein expression of haemoglobin is upregulated during monocyte differentiation and its levelis decreased as the differentiation progresses ^47–49^. Additionally, Giladi et al. ^38^, reports that *Itgb2, Itgam* is essential for neutrophil differentiation, which is consistent with CC-Tempo results. *Mmp8* and *Lyz2* are also reported as important in neutrophil differentiation, however, CC-Tempo found their expression important for monocyte. C*alr, Igfbp4*, important gene at progenitors for neutrophil as discussed above, continue to be important on the intermediate time points for neutrophil differentiation. Interestingly, some of the marker genes for monocyte and neutrophil (Supplementary Table 2) start expressing at intermediate time points; marker genes like *Ctss, Fabp5* and *Tgfbi* play critical role in monocyte differentiation at intermediate stages while *S100a8, S100a9* and *Lcn2* play critical role in neutrophil differentiation. Even though, *Fabp5* continues to be important on final time point, *Ctss* and *Tgfbi* plays no important role then for monocyte differentiation, while all of *S100a8, S100a9* and *Lcn2* continues to be important for neutrophil. Both *Fth1* and *Mrc1* has been reported as important for monocyte differentiation while *Gstm1* has been reported as important for Neutrophil differentiation by Weinreb et al. Both of these results are consistent with CC-Tempo output. However, *Cybb* and *Hspa5* has been reported as important for monocyte differentiation, while CC-Tempo finds them important for neutrophil. In addition, *Alas2, Vim, Lgals3, Psap, Ctsc* and *Isg15* all have been reported to play crucial role in monocyte differentiation ^43,48,50–53^ while *Tmsb4x, Wfdc21* and *Msrb1* have been reported to play important role in neutrophil differentiation ^54–56^. Finally, *Anxa5, Clec4n, Prdx5* and *Cd52* has no known role in monocyte neutrophil differentiation.

The important genes at the final time points, as reported by CC-Tempo, are also has strong experimental support in literature (Fig. 3A-B). Weinreb et al, reports that *Wfdc17* and *Fth1* are important for monocyte which is consistent with CC-Tempo results. Additionally, *Fabp5*, a marker gene for monocyte, has been reported as important as well while *S100a9, S100a8* and *Lcn2* continue to be important from intermediate time points. Giladi et al., reports that *S100a4* is important for monocyte differentiation while *Ngp, Fcer1g, Ltf,Ifitm6, Itgam* and *Syne1* is important for neutrophil differentiation. These results are consistent with CC-Tempo’s interpretation. However, *Lyz2*, an important gene for neutrophil has been reported as important for monocyte. *Psap* and *Lgals3* continue to affect the monocyte differentiation while *Timp2, Naaa, Ctsb, Gpnmb, Efhd2* come out to join the process at final time point. They have been reported to be important for monocyte differentiation in literature^57–61^. Additionally, *Wfdc21* and *Anxa1* has been reported as important for neutrophil differentiation ^55,62^, consistent with CC-Tempo’s result. *Mpeg1, Mcemp1, Cd33* and *Tyrobp* has no known role in monocyte neutrophil differentiation yet.

This sets strong ground for CC-Tempo’s interpretation results, supporting that CC-Tempo can consistently capture genes over the the cellular differentiation process that actually plays crucial role in the process.

### CC-Tempo can find cell-cell signaling pathways that determine cell fate bias and align with existing literature

CC-Tempo not only focuses on gene expression data but also the way cells communicate with each other to determine each other’s final fate. We used SHAP in the same fashion as above (See Sec. 4 (Methods)) to find the signaling pathways that were highly influential in determining the final cell fate.

For the Weinreb et al. dataset, we found out that 29 pathways are used for cell-cell communication using CellChat^19^ (See Sec. 4 (Methods)). Interestingly, for all the cell fate, CC-Tempo suggested that only two signaling pathways are highly important in determining cell fate, while most of the other pathways are mostly irrelevant. These pathways are *CD34* and *SELL. CD34* was important in sending signals, while *SELL* was used to receive signals in all cell types. CC-Tempo suggests that these pathways affect cell fate negatively for both monocytes and neutrophils, while for all other cells, these pathways affect cell fate positively (Fig. 6A).

**Figure 6:**
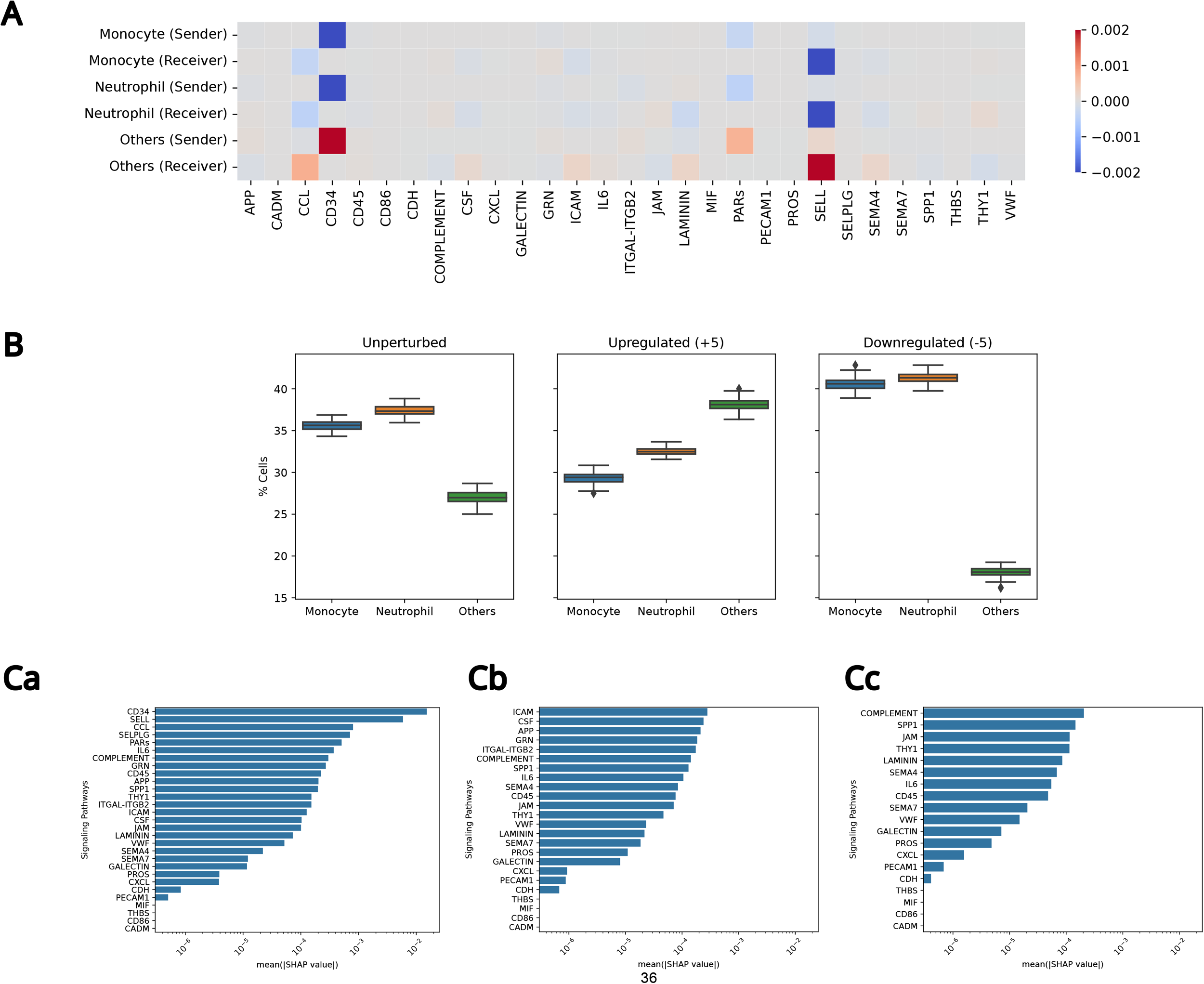
**A)** The importance of different pathways according to SHAP for determining the final day fates of cells. A cell can either send a signal via a specific pathway or receive a signal via that pathway, hence, it can be either sender or receiver or both. **B)** A boxplot showing the percentage of cells in different lineage when these important pathways, as well as their corresponding ligand and receptor, are perturbed in silico. **C)** Barplot of Mean Absolute SHAP values when CC-Tempo was trained on different numbers of signaling pathways. **a)** CC-Tempo was trained on all 29 pathways. **b)** CC-Tempo was trained on 24 pathways (Top 5 pathways were removed) c) CC-Tempo was trained on 19 pathways (Top 10 pathways were removed).

To validate this suggestion by CC-Tempo, we first calculate the percentage of monocyte, neutrophil, and other cells in the unperturbed settings, and then we perturb the cell-cell interaction score (+5) as well as the score of corresponding ligand and receptor (+5). When the perturbation is not introduced, the percentage of monocyte, neutrophil, and other cells were 35. 59 ± 0. 11, 37. 4 ± 0. 12, and 27. 01 ± 0. 14, respectively. When we upregulate the signaling pathways as well as the corresponding ligand-receptor expressions (+5), the number of other cells increases to 38. 13 ± 0. 13 while we downregulate the pathway and ligand-receptor to −5, the percentage of other cells decline to 18. 10 ± 0. 12 which fits exactly with the suggestions of CC-Tempo (Fig. 6B).

Such use of cell-cell interaction in determining cell fate has been largely ignored in the literature. However, CC-Tempo suggests a novel way to incorporate this vital information better to understand cellular developmental dynamics.

### CC-Tempo consistently orders signaling pathways in terms of importance without artificially inflating them, even when the most critical ones are excluded

Finally, we conducted an experimental evaluation of the role of signaling pathways in CC-Tempo’s ability to learn cellular dynamics. We assessed and ordered these pathways based on their significance by employing SHAP analysis, as shown in Figure 6A. This analysis identified the five key signaling pathways that play a pivotal role in determining the fate of monocytes, neutrophils, and other cell types: *CD34, SELL, CCL, SELPLG*, and *PARs* (Fig 6Ca). When CC-Tempo was trained without these key signaling pathways, the clonal fate bias metric decreased from 0.520 to 0.512, as shown in Figure 2D, indicating a slight performance reduction without these critical pathways. Upon reassessing the importance of the remaining pathways using SHAP analysis, the new top five critical pathways identified were *ICAM, CSF, APP, GRN*, and *ITGAL-ITGB2*, as depicted in Figure 6Cb. Notably, these pathways were already ranked within the top 15 for importance when CC-Tempo included all signaling pathways, as per Figure 6Ca. Despite the removal of the initially top-ranked pathways, these five signaling pathways did not increase in relative importance, confirmed by a Wilcoxon test (p-value > 0.01, see Supplementary Table 11). Furthermore, excluding the top 10 pathways revealed *COMPLEMENT, SPP1, JAM, THY1*, and *LAMININ* as the next five key pathways, shown in Figure 6Cc. These also were among the top 17 initially and maintained their levels of importance without any statistical significance in change (Wilcoxon test, p-value > 0.01, Supplementary Table 12). The exclusion of the top 10 pathways significantly impacted CC-Tempo’s efficacy, reducing the clonal fate bias metric from 0.52 to 0.50 (Fig. 2D). These findings suggest that CC-Tempo has a consistent methodology for ranking signaling pathways based on their contributions to cellular dynamics learning without artificially inflating the importance of any signaling pathways, even when some significant ones are absent.

## Discussion

Using only scRNA-seq data to comprehend the potential trajectory of cells can result in misidentification of their fate decisions. Approaches like Population Balance Analysis (PBA), WaddingtonOT, and FateID have been found to underperform compared to predictions based on the top ten genes^17^. Yeo et al. later demonstrated that incorporating a cell proliferation score improves the model’s ability to grasp the potential trajectory, leading to more accurate clonal fate determinations. Now, CC-Tempo demonstrates that leveraging inherent intercellular communication information within scRNA-seq data can significantly enhance decision accuracy. Importantly, CC-Tempo does not require any additional data beyond what previous methods use. It can extract intercellular communication scores directly from scRNA-seq data and, when integrated into the model, outperforms existing methods. CC-Tempo reveals that models relying solely on scRNA-seq data can be enhanced further by reusing scRNA-seq data from various dimensions. This suggests the possibility of future models that can predict cell fate decisions exclusively using scRNA-seq data, with CC-Tempo representing a step in that direction.

The highly variable genes (HVGs) in linear potential component only contains 26% genes that are important for intercellular communication, hence cannot recover intercellular communication accurately. When we added all the remaining intercellular communication specific genes to the set of HVGs and tried to recover the intercellular communication, CellChat inferred more than 50% false positive intercellular communication while only being able to capture <50% actual communication that were available in the dataset (Supplementary Table 13). Even though, most of the information in the intercellular component is indirectly available in the gene expression space, it would be hard for neural networks to capture these meaning intercellular communications from the exceedingly high dimensional space without any prior knowledge like ligand-receptor information from existing literature. Thus, with prior intercellular communication knowledge, the separate intercellular communication component guides the model in a more informed way to learn cellular dynamics, which is reflected in the performance of CC-Tempo. Integrating both of these components into a single model in a sensible and meaningful manner without hurting the model’s capacity to learn and interpret cellular dynamics would be an interesting future avenue for this work.

With its improved accuracy, CC-Tempo opens up new opportunities to hone a more accurate mechanistic understanding of how cells progress through differentiation or other temporal trajectories. Also, CC-Tempo prioritizes the genes and intercellular communication pathways whose perturbation can modulate the abundance of specific cell types. This can immediately help improve the efficiency of cellular differentiation protocols.

CC-Tempo’s second key feature lies in its interpretability. Many existing methods rely on low-dimensional principal components (PCs) of scRNA gene expression space, making it challenging to discern which genes influence cellular trajectories at various stages. In contrast, CC-Tempo uses the raw scRNA-seq data as its input, enabling it to be interpreted using various neural network explanation techniques. CC-Tempo introduces a method for elucidating the outcomes of cellular dynamics models and ranking the genes and pathways that play a crucial role in guiding cells toward their ultimate fate. This interpretative framework can be applied to any neural network-based model, extending beyond CC-Tempo itself. This development opens up a novel avenue for comprehending the functioning of diverse neural networks in the realm of biology.

Looking ahead, CC-Tempo holds promise for applications in understanding various cellular dynamics processes and disease progression. There is an exciting potential for incorporating additional data modalities, such as spatiotemporal data and epigenetics data, in future research directions.

## Methods

### Learning cellular differentiation dynamics through diffusion

Recent studies have used diffusion processes to model cell state evolution in single-cell populations^18,31^. However, prior models have focused exclusively on the evolution of transcriptomic states (gene expression values). We propose a model, CC-Tempo (A cell-cell communication aware temporal model of cellular dynamics), where we incorporate cell-cell communication in modeling cellular differentiation and cell state evolution. In particular, we model cellular differentiation as a tuple of differentiation processes ((*X*(*t*), *C*(*t*)) represented by the following stochastic differential equations

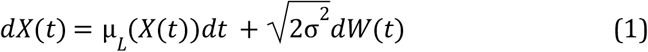

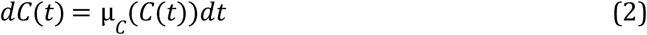

where *X*(*t*) represents the *k*-dimensional gene expression component of the state of a cell at time point *t* while *C*(*t*) represents *l*-dimensional intercellular communication score component of the state of a cell at time point *t*. μ_*L*_ (.) and μ_*C*_ (.) are two functions representing drift velocities: μ_*L*_(*X*(*t*)) represents the drift velocity of the cell in linear potential space due to force acting upon the cell to move from higher to lower potential. Similarly, μ_*C*_(*C*(*t*)) denotes the drift velocity of the cell in the intercellular communication potential space, and finally *W*(*t*) is the Wiener process that adds some noise. This noise is only added to the linear potential space rather than in intercellular communication space because adding it to the later space didn’t improve performance any further. Both of the drift velocities are negative gradients of their corresponding potential function, i.e.,

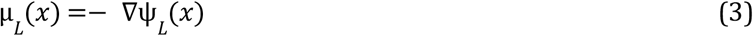

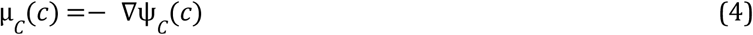

These potentials invoke a gradient field that drives cells from higher potential to lower potential regions in their corresponding space. We can simulate this process via a first-order time discretized equation as follows

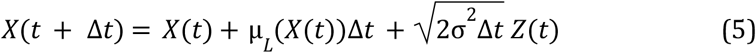

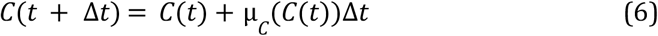

where *Z*(*t*) are i.i.d standard Gaussian noise, *N*(0, 1). These equations will converge to the diffusion process when Δ*t* → 0.

We define two marginal probability distributions at a particular time point t which are ρ_*L*_(*x, t*) = *P*_*L*_(*X*(*t*) = *x*) and ρ_*C*_(*c, t*) = *P*_*C*_(*C*(*t*) = *c*). In particular, we suppose that we are given some observed samples from these distributions at a few particular snapshots of time, which are {*x*(*t*)_*i*_ ∼ ρ_*L*_(*x, t*)|*i* ∈ {1 … *m*_*t*_}, *t* ∈ {1 … *n*}} and {*c*(*t*)_*i*_ ∼ ρ_*C*_(*c, t*)|*i* ∈ {1 … *m*_*t*_}, *t* ∈ {1 … *n*}} where *m*_*t*_ is the number of cells sampled at the time point *t* and *n* is the numbers of timepoints where data were observed, *x*(*t*)_*i*_ is obtained directly from gene expression data while *c*(*t*)_*i*_ is obtained from the intercellular communication module using the gene expression as input. Now, we try to learn the potential functions ψ_*L*_(*x*) and ψ_*C*_(*c*) which in turn allows us to learn the corresponding drift velocity via simulating these samples in the above first-order time discretized equation and the corresponding marginal probability distribution.

The learning proceeds by deriving the set of potential functions in the families of function space (*K, M*) that minimizes the objective loss function

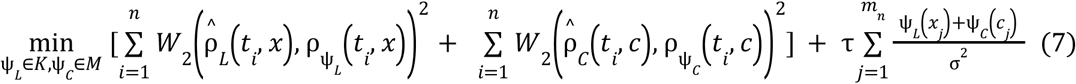

where 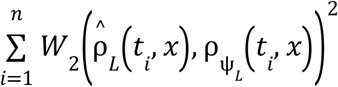 is the Wasserstein distance between the empirical distribution 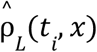 obtained from observed gene expression data and the distribution of the candidate potential function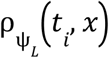. Similarly, 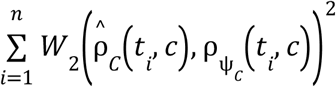is the Wasserstein distance between the empirical distribution 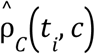 obtained from the observed intercellular communication data and the distribution of the candidate potential function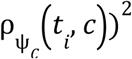. τ is the parameter that controls the strength of the entropic regularizer term. Wasserstein loss function has been widely used in literature for such optimization ^63^.

### Incorporating Cellular Proliferation

Following the notation of Feydy et al.^64^, we define the optimization problem using Wasserstein distance as follows

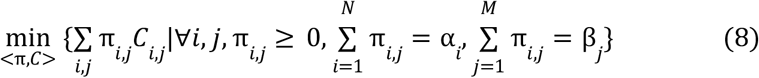

where π _*i,j*_ is the optimal transport plan that maps points from source distribution to the target distribution and *C*_*i,j*_ =‖ *x*_*i*_ − *y*_*j*_ ‖^2^ is the square of the Euclidean distance between the sample *x*_*i*_ and sample *y*_*j*_, and finally α_*i*_ and β_*j*_are positive weights associated with each sample *i* and *j*. Following Yeo et al., we incorporated cellular proliferation into CC-Tempo by setting α_*i*_ to the number of descendant cells the sample *i* is expected to have and keep β_*j*_ constant. In particular, we estimate the number of descendants in the following manner. Let *n* denote the number of descendants of the sample *i, b* denote the birth rate, *d* denote the death rate, and finally *g* denote the growth rate. Following the birth-death process, for a given clone,

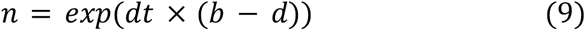

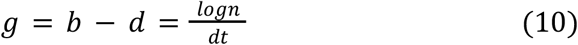

To calculate the birth score *s*_*b*_, we calculated the mean of z-scores of the genes annotated to the cell cycle pathway *(KEGG Cell Cycle)*. Similarly, to calculate, *s*_*d*_, we calculated the mean z-scores of the genes annotated to the cell death pathway (*KEGG Apoptosis)*. Then these scores were smoothed over the cells using the following iterative procedure:

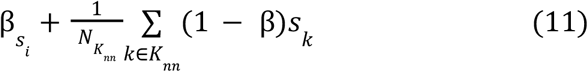

where *s*_*i*_ is the score for a sample i at the current iteration while *s*_*k*_ is the score for *k* nearest neighbors of the cell i.

Finally, the birth and death scores are obtained using the following equations

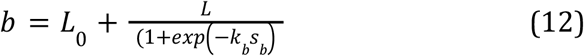

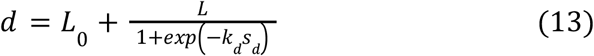

We set the value of the hyperparameters using the values proposed by Yeo et al. Note, however, that we did not incorporate the same in our cell-cell interaction potential function learning as it did not improve our performance any further.

### Running CellChat to infer Signaling Pathways Score

We use CellChat to find out the enriched signaling pathways in our desired dataset, In this case, mouse hematopoesis data by Weinreb et al. We used the traditional pipeline described in the CellChat documentation with no parameters tweaking. The following methods were run sequentially to obtain the score for signaling pathways.

~~~
cellchat <- subsetData(cellchat)
cellchat <- identifyOverExpressedGenes(cellchat)
cellchat <- identifyOverExpressedInteractions(cellchat)
cellchat <- computeCommunProb(cellchat)
cellchat <- filterCommunication(cellchat, min.cells = 10)
cellchat <- computeCommunProbPathway(cellchat)
cellchat <- aggregateNet(cellchat)
~~~

### Incorporating Intercellular Communication Score

First, we use CellChat to calculate the incoming communication and outgoing communication matrix *I* and *O* respectively. Each entry 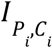 of the incoming communication matrix *I* ∈ *R* ^*C*×*P*^ denotes the probability scores of cell type *C*_*i*_ receiving an intercellular signal via pathway *P*_*i*_. In a similar manner, each entry 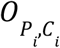 of the outgoing communication matrix *O* ∈ *R* ^*C*×*P*^ denotes the probability scores of cell type *C*_*i*_ sending an intercellular signal via pathway *P*_*i*_. Each pathway is defined in terms of its ligand-receptor pairs, i.e., it has a set of ligands 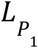 and a set of receptors 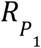. So, initially, we calculate the ligand score 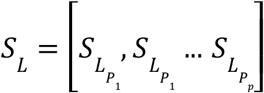 and the receptor score 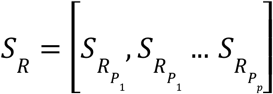 of each cell via the *Seurat*^65^ function *AddModuleScore*. Then using the cell type information, we take the Hadamard product corresponding row of the incoming matrix *I* and receptor score *S*_*R*_ to obtain our receiving score for the corresponding cell.

Similarly, we take the Hadamard product corresponding row of the outgoing matrix *O* and ligand score *S*_*L*_ to obtain our sending score. Finally, we concatenate these two scores, the sending score and receiving score to get the input of our intercellular communication network, i,e. intercellular communication score (Fig. 1A).

#### Model Definition

We followed a modularized implementation of CC-Tempo and built different components of CC-Tempo in part. We describe each of them in the following points.

- **Potential Learning Network**. We have two feed-forward neural networks to learn the linear potential and intercellular communication potential, respectively. Both of them have two layers where the linear potential network takes the low dimensional representation of gene expression as its input and outputs a scalar linear potential value. In contrast, the intercellular communication network takes the concatenated vector of the sending and receiving score as derived above as input and outputs a scalar intercellular communication potential. Finally, we take the sum of these potentials and output the total potential (Fig. 1Bi).
- **Gradient Calculator** To calculate the drift velocity from potential, we need the gradient of our corresponding potential function. So, in this network, we pass the potential score provided by our Potential Learning Network and calculate the gradient with respect to our two inputs *x* and *c*. Thus, the outputs of this network are two gradient scores Δ_*x*_ and Δ_*c*_ respectively which are our drift velocities (Fig. 1Bii).
- **SHAP** For calculating the SHAP score, two implementations from the Captum^66^ library were used, one is GradientSHAP, and another is KernelSHAP. Both of them are two different implementations to calculate SHAP scores and require the corresponding model to be implemented in the PyTorch framework (Fig. 1Biii). captum.attr.**GradientShap(model)**.attribute(*inputs, baselines, n_samples=5, stdevs=0*.*0, target=None, additional_forward_args=None, return_convergence_delta=False*) captum.attr.**KernelShap(model)**.attribute(*inputs, baselines=None, target=None, additional_forward_args=None, feature_mask=None, n_samples=25, perturbations_per_eval=1, return_input_shape=True, show_progress=False*)
- **PCA and inverse-PCA** We built a feed-forward neural network to implement the Principal Component Analysis. First, we calculated the PCA of high-dimensional gene expression using *scikit-learn*^*67*^, which reduced the high-dimensional input gene expression into Low-dimensional PCs. Then we created a neural network whose input is High-dimensional input gene expression and output is Low-dimensional PCs. *scikit-learn* provides a matrix of dimension *R* ^*gene*×*PC*^, which we set as the weight of our neural network instead of training the network. We need our dimension reduction model to be a neural network in order to evaluate it using SHAP, as we describe later. Hence, we need to build PCA as such. Similarly, we can transpose the matrix from *scikit-learn* and build another feed-forward neural network that converts the low-dimensional PC input into high-dimensional gene expression input (Fig. 1Biv). *class* sklearn.decomposition.**PCA**(*n_components=50, *, copy=True, whiten=False, svd_solver=‘auto’, tol=0*.*0, iterated_power=‘auto’, n_oversamples=10, power_iteration_normalizer=‘auto’, random_state=None*)
- **Cell Type Classifier** We use logistic regression as our classifier to compare CC-Tempo with PRESCIENT so that the performance improvement is solely based on the model itself, not on the strength of the classifier. But for SHAP, we need CC-Tempo to be a fully neural network. So, where CC-Tempo is required to be a fully neural network, we used a two layer neural network to classify our cells (Fig. 1Av). *class* sklearn.linear_model.**LogisticRegression**(*penalty=‘l2’, *, dual=False, tol=0*.*0001, C=1*.*0, fit_intercept=True, intercept_scaling=1, class_weight=None, random_state=None, solver=‘lbfgs’, max_iter=100, multi_class=‘auto’, verbose=0, warm_start=False, n_jobs=None, l1_ratio=None*)
- **CC-Tempo**. Finally, CC-Tempo takes a high-dimensional gene expression as its input. Then it passes a copy of this expression to the Cell-Cell Interaction Module to get the intercellular communication score which is used as an input to the intercellular communication potential network. while we pass the first copy of gene expression through our PCA adapter and get the low dimensional representation and pass it to our linear potential network. Then, we get the output of both of these potential networks, sum them, and pass it to our gradient calculator to get the drift velocity. Then we use (Equation 1) to simulate the cell to the next time point and continue simulating it till the last time point, where we again use the inverse-PCA adapter to convert the cell back into gene expression space, and this is the final output of CC-Tempo (Fig. 1C).

#### Model Optimization

Both of our potential learning networks have identical structures with different input dimensions. In particular, the linear potential ψ_*L*_(*x*) takes 50-dimensional input, which is the output of our PCA Adapter, and has two fully connected layers with 500 units and 1 unit, respectively. On the other hand, the intercellular communication potential ψ_*C*_(*c*) takes the concatenated input of sending and receiving scores which is 29 × 2 = 58 dimensional and has the same two fully connected layers as the architecture. We used PyTorch^68^ as our framework, and we used PyTorch’s automatic differentiation to derive our corresponding drift velocity. We set our time step *dt* to be 0. 1 while the entropic regularizer parameter τ to be 1*e* − 6. On each iteration, we took 10% of the training data and simulated through the network to compute cost and backpropagate. We used Adam Optimizer to optimize CC-Tempo. We used the pretraining methods of the model described by Hashimoto et al. to pre-train CC-Tempo before the final training stage. The Wasserstein distance was calculated by the Sinkhorn algorithm implemented by the GeomLoss library^64^, which is compatible with PyTorch GPU tensors. The model was pre-trained for 500 epochs while it was trained for 2500 epochs.

All of the models were trained in Google Colab Pro+ with a single NVIDIA A100 40 GB GPU and 81 GB memory. On average, training the full network on the full Weinreb et al. dataset took ∼ 30*mins*.

#### Predicting Clonal Fate bias

Clonal fate bias is the metric defined by Weinreb et al. to evaluate the performance of the model in determining cell fate. It is calculated as the number of neutrophils divided by the total number of neutrophils and monocytes in the particular clone. Following Prasad et al.^69^, we extract only those Day 2 progenitor cells that have at least one neutrophil or monocyte in their Day 4 or Day 6 fate and use these cells to validate CC-Tempo (Fig. 2B). We simulated these progenitor cells via our trained model till Day 6. Then we used the trained logistic regression classifier to classify the cell into monocyte, neutrophil, or other cell types. In particular, we made 2000 copies of each of our progenitor cells and simulated via CC-Tempo to get the final state of the cells. Due to the random component in CC-Tempo, 2,000 cells went into 2,000 different states. Then we classified these 2000 cells again into monocyte, neutrophil, or other cells via our logistic regression classifier. Finally, we calculated the clonal fate bias of that cell as the number of neutrophils divided by the total number of neutrophils and monocytes.

Since CC-Tempo did not always predict neutrophils or monocytes as the cell fate, we added a pseudocount of 1 to avoid division by zero, and hence in that case, our clonal fate bias score will be 0. 5. Finally, we calculated the PCC and AUROC score between the CC-Tempo predicted value and values scored from lineage tracing data to benchmark CC-Tempo (Fig. 1Di).

#### Benchmarking with PRESCIENT

We used the default implementation of PRESCIENT, provided with paper without tweaking any default parameters.

~~~
prescient train_model *-i data*.*pt --out_dir /experiments/ --weight_name
‘kegg-growth’ --seed 2*
~~~

#### Clonal Fate Deviation

We define a new metric for clonal fate deviation to calculate how close the model’s predicted fate bias is to actual fate bias. To do so, we take the absolute value of the difference between the model-predicted bias and actual fate bias and divide it via actual fate bias.

#### Wilcoxon’s rank-sum Test

We used Python’s *Scipy* libraries implementation of Wilcoxon’s rank-sum test. All the tests were conducted using 5% statistical significance.

scipy.stats.**ranksums**(*x, y, alternative=‘two-sided’, *, axis=0, nan_policy=‘propagate’, keepdims=False*)

#### Obtaining cell-fate-defining genes

We used SHAP values to determine the potential cell-fate-defining genes in different cell types. We took all the progenitor cells in our dataset and simulated through CC-Tempo to get the Day 6 cell state. Then, we classified the cells using our trained logistic regression to classify them into neutrophils, monocytes, or other cells. Then, we took the final day states (calculated by CC-Tempo) of the cells that were predicted monocytes and then used the SHAP score to determine which genes were responsible for determining their fate as monocytes. In particular, the inputs of our SHAP model are the final day cell state predicted by CC-Tempo, and the output is the total potential value. We determined the inputs(genes) that were responsible for taking those cells into that particular potential score on the final day. In a similar fashion, we determined the cell-fate-defining genes for neutrophils as well (Fig. 1Dii).

#### Transcription Factor’s Perturbation and Evaluation

We have already obtained transcription factors for neutrophils and monocytes from the literature. We manually set the value of these transcription factors in the dataset and re-simulated the cells via CC-Tempo to check whether the introduction of progenitor cell’s transcription factors’ perturbation changes the trajectory of progenitor cells. To upregulate a transcription factor, we gradually increased the expression value for that gene from the initial value to 1, 2, 2. 5. Then we simulated all the progenitor cells 100 times to get the percentage of monocytes or neutrophils and plotted them with 95% Confidence Interval. Similarly, when downregulating a transcription factor, we decreased the transcription factor’s value from the initial value to − 1, − 2, − 2. 5 and followed similar steps.

#### Obtaining cell fate-defining genes in progenitor cells and perturbation evaluation

We then tried to predict whether CC-Tempo can determine the progenitor cells’ genes that were influential in the determination of the final day cell fate. In Particular, we took the progenitor cells whose fate were monocytes(neutrophils) and simulated through CC-Tempo to get the Day 6 cell state and then the potential value from that state. We used SHAP to evaluate the importance of progenitor cells’ genes importance in determining final day potential value. Specifically, the input of our SHAP model is the Day 2 progenitor cells, and the output of CC-Tempo is the Day 6 potential value. In this way, we obtained the genes that are important for the Day 6 cell fate determination (Fig. 1Diii).

#### Obtaining Relevant Signaling Pathway

We also focused on determining different signaling pathways’ importance in progenitor cells that determine the final day cell fate. Like above, we again used SHAP to order our signaling pathway according to their importance in determining the corresponding cell fate. Particularly, our input now is the intercellular communication score of Day 2 progenitor cells, and the output is the cell fate, i.e., whether the cell is monocyte, neutrophil, or other types of cell. Then we sorted the pathways according to their importance and found the relevant pathways for each cell type (Fig. 1Div).

#### Perturbing Signaling Pathway and Evaluation

Finally, we validated if the signaling pathways determined by CC-Tempo can actually change cell fate. So, we set the intercellular communication of the pathway as well as the gene expression of the ligand-receptor pairs of that pathway to + 5 upregulate the pathway and − 5 to downregulate the signaling pathway. Finally, we simulated each perturbed configuration 100 times to get the percentage of cell type in those configurations.

## Code & Data Availability

All the code, from data preprocessing to model evaluation, can be found at https://github.com/Srj/CC-Tempo. The raw LT-scSeq mouse hematopoiesis dataset was obtained from https://github.com/AllonKleinLab/paper-data/tree/master/Lineage_tracing_on_transcriptional_landscapes_links_state_to_fate_during_differentiation.

