## Supplementary Figure for "CC-Tempo: A cell-cell communication aware temporal model of cellular dynamics"

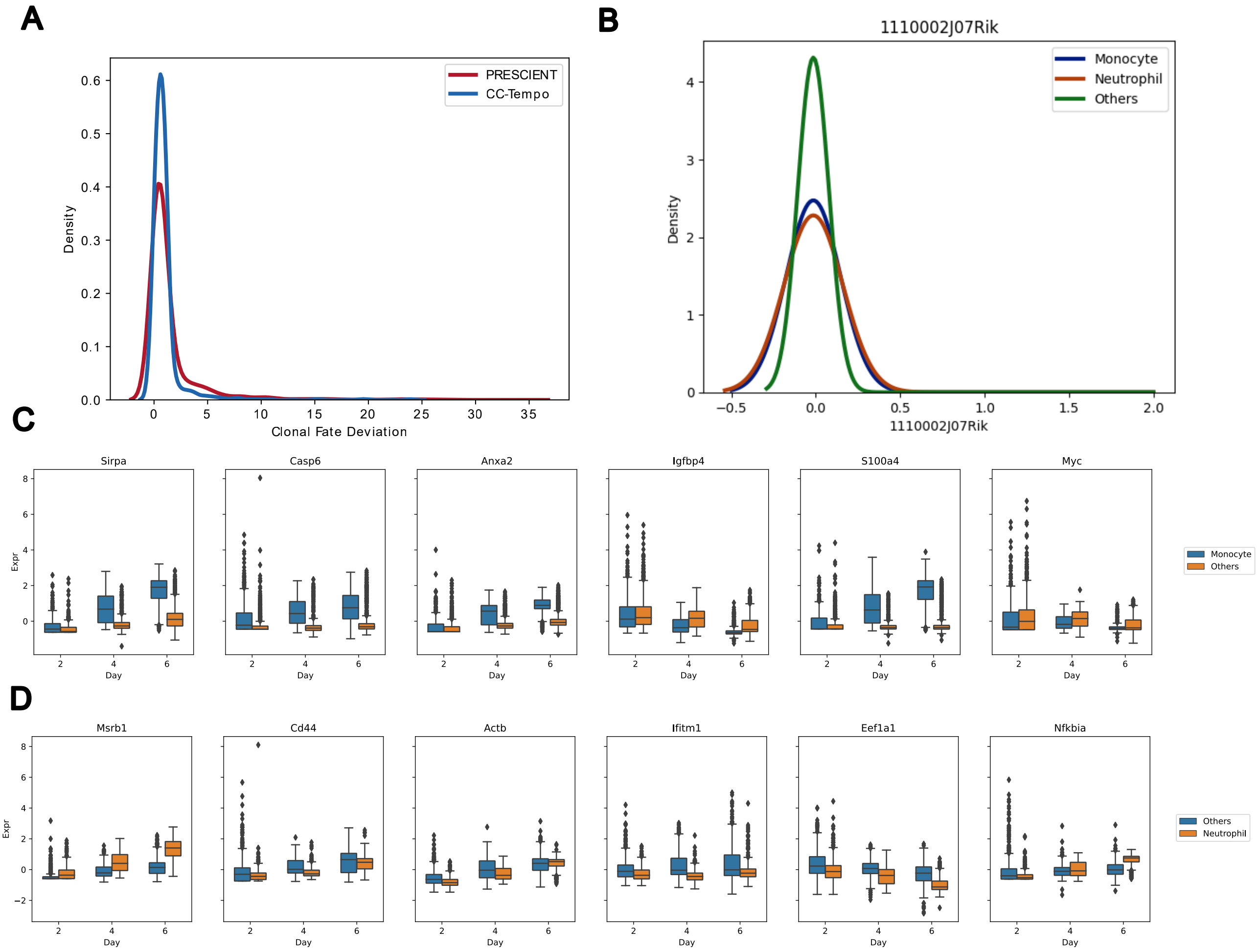

A
PRESCIENT
CC-Tempo
Density
Clonal Fate Deviation
B
1110002J07Rik
Monocyte
Neutrophil
Others
Density
1110002J07Rik
C
Sirpa
Casp6
Anxa2
Igfbp4
S100a4
Myc
Expr
Day
Monocyte
Others
D
Msrb1
Cd44
Actb
Ifitm1
Eef1a1
Nfkbia
Expr
Day
Others
Neutrophil

**Supplementary Figure 1:**

**A)** The distribution plot of clonal fate deviation for PRESCIENT and CC-Tempo. **B**) The distribution plot of the gene *1110002J07Rik* for neutrophil, monocyte, and other cell types. **C)** The gene expression plot for the genes that are important for monocytes on all time points (Day 2,4,6) **D)** The gene expression plot for the genes that are important for neutrophils on all time points (Day 2,4,6)
