## Supplementary Table for "CC-Tempo: A cell-cell communication aware temporal model of cellular dynamics"

**Supplementary Table 1**

| Alternative Hypothesis | p-value |
|---|---|
| CC-Tempo’s score (Clonal Fate Bias) is greater than PRESCIENT | 7e-19 |
| CC-Tempo’s score (Clonal Fate Bias) is greater than CC-Tempo w/ Top 5 pathways removed | 0.09 |
| CC-Tempo’s score (Clonal Fate Bias) is greater than CC-Tempo w/ Top 10 pathways removed | 7e-5 |
| CC-Tempo w/ Top 5 pathways removed’s score is greater than CC-Tempo w/ Top 10 pathways removed | 0.02 |
| CC-Tempo w/ Top 5 pathways removed’s score is greater than PRESCIENT | 0.001 |
| CC-Tempo w/ Top 10 pathways removed’s score is greater than PRESCIENT | 0.52 |

**Supplementary Table 2**

| Cell type | Marker Gene | Reference |
|---|---|---|
| Monocyte | Ms4a6d | http://journals.plos.org/plosone/article?id=10.1371/journal.pone.0164027 |
| | Fabp5 | https://www.ncbi.nlm.nih.gov/pubmed/26105806 |
| | Ctss | https://www.ncbi.nlm.nih.gov/pubmed/16365419 |
| | Ms4a6c | http://journals.plos.org/plosone/article?id=10.1371/journal.pone.0164027 |
| | Tgfbi | http://journals.plos.org/plosone/article?id=10.1371/journal.pone.0072772 |
| | Olfm1 | https://www.nature.com/articles/s41698-017-0031-0/tables/1 |
| | Csf1r | https://www.ncbi.nlm.nih.gov/pmc/articles/PMC5009875/ |
| | Ccr2 | https://www.nature.com/articles/nri2999 |
| | Klf4 | https://www.ncbi.nlm.nih.gov/pmc/articles/PMC2230668/ |
| | F13a1 | http://www.jimmunol.org/content/177/10/7303 |
| Neutrophil | S100a9 | http://www.jimmunol.org/content/jimmunol/170/6/3233.full.pdf |
| | Itgb2l | https://www.ncbi.nlm.nih.gov/pubmed/15922663 |
| | Elane | |
| | Fcnb | https://www.ncbi.nlm.nih.gov/pubmed/22459270 |
| | Mpo | http://www.bloodjournal.org/content/117/3/953 |
| | Prtn3 | https://www.ncbi.nlm.nih.gov/pubmed/9925946 |
| | S100a6 | Neutrophils, The: New Outlook For Old Cells (3rd Edition), page 312 |
| | S100a8 | Neutrophils, The: New Outlook For Old Cells (3rd Edition), page 312 |
| | Lcn2 | https://www.ncbi.nlm.nih.gov/pubmed/22965758 |
| | Lrg1 | https://journals.plos.org/plosone/article?id=10.1371/journal.pone.0170261 |

**Supplementary Table 3**

| Gene | p-value |
|---|---|
| 1110002J07Rik | 0.987632 |
| Mrc1 | 0 |
| Wfdc17 | 0 |
| Timp2 | 0 |
| Mpeg1 | 0 |
| S100a4 | 0 |
| Psap | 0 |
| Naaa | 0 |
| Fabp5 | 0 |
| Cstb | 0 |
| Lgals3 | 0 |
| Gpnmb | 0 |
| Fth1 | 0 |
| Lyz2 | 0 |
| Efhd2 | 0 |

**Supplementary Table 4**

| **Gene** | **p-value** |
| --- | --- |
| 1110002J07Rik | 0.9829583 |
| Ngp | 0 |
| Lcn2 | 0 |
| Mcemp1 | 0 |
| S100a9 | 0 |
| Fcer1g | 3.56E-300 |
| Wfdc21 | 0 |
| Ltf | 0 |
| Ifitm6 | 0 |
| Itgam | 0 |
| Cd33 | 0 |
| S100a8 | 0 |
| Syne1 | 0 |
| Tyrobp | 0 |
| Anxa1 | 0 |

### Supplementary Table 5

| Gene | p-value |
|---|---|
| Igfbp4 | 0.0291525578977226 |
| Casp6 | 5.36E-24 |
| Samhd1 | 0.00425424763311525 |
| Ifi203 | 1.16E-22 |
| Prdx1 | 4.12E-09 |
| Muc13 | 5.09E-18 |
| Rbms1 | 3.18E-14 |
| Spint2 | 5.98E-06 |
| Srgn | 1.64E-35 |
| Tpd52 | 2.32E-06 |
| Atp1b3 | 0.0062856717334331 |
| Myc | 0.000580667718794982 |
| Calr | 5.33E-22 |
| H2-Q7 | 0.000168079385354811 |
| Pou2f2 | 2.34E-15 |

### Supplementary Table 6

| Gene | p-value |
|---|---|
| Srgn | 3.49E-59 |
| Tmed3 | 9.77E-33 |
| Calr | 2.20E-59 |
| Gstm1 | 9.78E-56 |
| Igfbp4 | 0.76179834377803 |
| Dstn | 7.07E-50 |
| Muc13 | 0.0477587744973364 |
| Spint2 | 0.0032910322710516 |
| Prdx1 | 1.42E-05 |
| Ifi203 | 2.86E-19 |
| Igf1r | 7.83E-28 |
| Rbms1 | 0.925336838394158 |
| Gsr | 5.20E-56 |
| Cd34 | 3.10E-08 |
| Myc | 0.00471632925605433 |

**Supplementary Table 7**

| Gene | p-value |
|---|---|
| Hba-a2 | 3.34E-09 |
| Hbb-bs | 5.08E-09 |
| Hba-a1 | 3.08E-09 |
| Slc4a1 | 1.51E-10 |
| Alas2 | 3.77E-08 |
| Hbb-bt | 0.0173605554819343 |
| Vim | 3.04E-111 |
| Lgals3 | 2.65E-82 |
| Ctss | 1.20E-86 |
| Mmp8 | 6.47E-131 |
| Anxa5 | 3.54E-65 |
| Psap | 6.45E-130 |
| Fth1 | 4.10E-147 |
| Ctsc | 2.04E-160 |
| Lyz2 | 2.34E-55 |
| Fabp5 | 3.52E-109 |
| Clec4n | 5.79E-107 |
| Tgfbi | 2.09E-139 |
| Mrc1 | 7.37E-110 |
| Isg15 | 1.62E-31 |

### Supplementary Table 8

| Gene | p-value |
|---|---|
| Hba-a2 | 0.0220990444988353 |
| Hbb-bs | 0.154989681257536 |
| Hba-a1 | 0.028873295378841 |
| Slc4a1 | 0.00126148577080285 |
| Alas2 | 0.0099895081922688 |
| Tmsb4x | 2.41E-40 |
| Itgb2 | 2.92E-17 |
| Calr | 0.0024166259064848 |
| Itgam | 6.23E-19 |
| S100a9 | 5.16E-65 |
| Gstm1 | 8.96E-183 |
| S100a8 | 1.91E-66 |
| Wfdc21 | 8.19E-40 |
| Cybb | 0.00515402819662233 |
| Prdx5 | 1.66E-120 |
| Igfbp4 | 0.355427427158001 |
| Lcn2 | 2.95E-29 |
| Cd52 | 0.11705270260677 |
| Hspa5 | 1.79E-12 |
| Msrb1 | 3.52E-78 |

### Supplementary Table 9

| Gene | Day | p-value |
|---|---|---|
| Anxa2 | 2 | 0.257206719096133 |
| Anxa2 | 4 | 1.42E-73 |
| Anxa2 | 6 | 2.74E-120 |
| Sirpa | 2 | 2.56E-08 |
| Sirpa | 4 | 3.26E-108 |
| Sirpa | 6 | 5.30E-144 |
| S100a4 | 2 | 6.60E-08 |
| S100a4 | 4 | 1.10E-142 |
| S100a4 | 6 | 4.86E-167 |
| Igfbp4 | 2 | 0.0291525578977226 |
| Igfbp4 | 4 | 1.63E-48 |
| Igfbp4 | 6 | 5.41E-61 |
| Casp6 | 2 | 5.36E-24 |
| Casp6 | 4 | 1.78E-131 |
| Casp6 | 6 | 4.11E-98 |
| Myc | 2 | 0.000580667718794982 |
| Myc | 4 | 5.66E-19 |
| Myc | 6 | 0.000179385923276647 |

### Supplementary Table 10

| Gene | Day | p-value |
|---|---|---|
| Eef1a1 | 2 | 4.10E-22 |
| Eef1a1 | 4 | 5.98E-55 |
| Eef1a1 | 6 | 1.62E-93 |
| Actb | 2 | 5.46E-30 |
| Actb | 4 | 2.92E-21 |
| Actb | 6 | 0.015224613331127 |
| Ifitm1 | 2 | 1.12E-16 |
| Ifitm1 | 4 | 5.16E-74 |
| Ifitm1 | 6 | 2.14E-14 |
| Nfkbia | 2 | 7.02E-08 |
| Nfkbia | 4 | 0.00298921555272763 |
| Nfkbia | 6 | 5.29E-103 |
| Msrb1 | 2 | 2.99E-31 |
| Msrb1 | 4 | 1.45E-81 |
| Msrb1 | 6 | 7.14E-147 |
| Cd44 | 2 | 1.00E-08 |
| Cd44 | 4 | 5.08E-51 |
| Cd44 | 6 | 0.000114171164116901 |

### Supplementary Table 11

| Signaling Pathways | p-value |
|---|---|
| ICAM | 0.59298 |
| CSF | 0.6875 |
| APP | 0.6875 |
| GRN | 0.6875 |
| ITGAL-ITGB2 | 0.59298 |

### Supplementary Table 12

| Signaling Pathways | p-value |
|---|---|
| COMPLEMENT | 0.6875 |
| SPP1 | 0.21875 |
| JAM | 0.03125 |
| THY1 | 0.6875 |
| LAMININ | 0.5625 |

**Supplementary Table 13**

| **Particulars** | **Value** |
|---|---|
| Day 2 | |
| Number of CellChat Gene Inferred from Inverse PCA GEX | 2000 |
| Number of CellChat Gene Inferred from Actual Data | 953 |
| Length of Intersection | 494 |
| Day 4 | |
| Number of CellChat Gene Inferred from Inverse PCA GEX | 1992 |
| Number of CellChat Gene Inferred from Actual Data | 947 |
| Length of Intersection | 554 |
| Day 6 | |
| Number of CellChat Gene Inferred from Inverse PCA GEX | 1942 |
| Number of CellChat Gene Inferred from Actual Data | 958 |
| Length of Intersection | 571 |
